# Coastal bacteria and protists assimilate viral carbon and nitrogen

**DOI:** 10.1101/2024.06.14.598912

**Authors:** Joaquín Martínez Martínez, David Talmy, Jeffrey A. Kimbrel, Peter K. Weber, Xavier Mayali

## Abstract

Free viruses are the most abundant type of biological particles in the biosphere, but the lack of quantitative knowledge about their consumption by heterotrophic protists and bacterial degradation has hindered the inclusion of virovory in biogeochemical models. Here, using isotope-labeled viruses added to three independent microcosm experiments with natural microbial communities followed by isotope measurements with single-cell resolution and flow cytometry, we quantified the flux of viral C and N into virovorous protists and bacteria and compared the loss of viruses due to abiotic vs biotic factors. We found that some protists can obtain most of their C and N requirements from viral particles and that viral C and N get incorporated into bacterial biomass. We found that bacteria and protists were responsible for increasing the daily removal rate of viruses by 33% to 85%, respectively, compared to abiotic processes alone. Our laboratory incubation experiments showed that abiotic processes removed roughly 50% of the viruses within a week, and adding biotic processes led to a removal of 83% to 91%. Our data provide direct evidence for the transfer of viral C and N back into the microbial loop through protist grazing and bacterial breakdown, representing a globally significant flux that needs to be investigated further to better understand and predictably model the C and N cycles of the hydrosphere.

## Introduction

Viruses are an abundant, diverse, and dynamic component of marine systems, where they play a multitrophic role in the cycling of marine organic matter, but the fate of aquatic viral particles is understudied. The bulk of research on aquatic viruses has focused on microbial lytic infections in which a virus enters a susceptible and permissive host and hijacks the host’s replication machinery resulting in relatively quick cell lysis and the release of virus particles— virions—and cellular organic material. Viruses are estimated to kill 20-30% of the ocean’s single-celled organisms daily [1–3]. It is on this premise that viruses have been described as modulators of microbial community dynamics and productivity [1, 4–6] and catalysts for global biogeochemical cycling by shifting cellular organic matter away from higher trophic levels into the dissolved fraction, a process known as the viral shunt [7, 8]. Furthermore, viral lysis has more complex effects than first envisaged; for example, its role in carbon (C) export to the deep ocean is receiving increased attention. Viral lysis stimulates phytoplankton and bacterial production of transparent exopolymeric particles (TEP) that aggregate with cell debris, living microbes, and fecal pellets to form marine snow enhancing C flow through the biological pump, a process known as the viral shuttle [9–13]. A less studied aspect than the viral shunt and shuttle is the direct role of viruses on the trophic transfer of nutrients and energy up the food web, either through differential ingestion of infected prey [14, 15] or direct consumption of virions by phagotrophic protists, a process known as virovory [16–21]. The magnitude of each of these three virally-mediated processes of regeneration, sequestration, and export up the food web (shunt, shuttle, virovory) remains largely unresolved.

Virus particles are the most numerous biological entities in the ocean. Based on concentrations typically ranging from 10^6^ to 10^8^ virions mL^-1^ across ocean environments [22] and assuming 0.05– 0.2 fg C per virion [23, 24], the standing stock of virus particles in the global ocean may translate into 30–200 Mton C [25, 26]. The ‘viral shunt’ is estimated to release 145 Gton C year^-1^ (average 400 Mton C day^-1^) into the dissolved organic pool [27], which is operationally defined as the fraction passing through a 0.2 – 0.7 µm filter pore size [28] and thus includes a majority of virus particles and small bacteria. In addition to C, virions comprise nitrogen (N) and phosphorus (P) as they are made up of proteins, nucleic acids, and lipids (in some taxa) [24], and even micronutrients [29, 30]. Consequently, virion consumption may be a significant marine microbial food web pathway that redirects nutrients and energy to upper trophic levels.

Over the last few decades, many studies have addressed the viral shunt and shuttle, but we lack quantitative data on the flow of nutrients and energy mediated by virovory across marine habitats and seasons to parameterize food web and biogeochemical models. Notably, virtually nothing is known about elemental transfer from viruses to bacteria. Regarding virovory by protists, an early study with natural coastal communities and enrichment cultures of nanoflagellates, the most important grazers of bacteria across many aquatic ecosystems [31], demonstrated phagotrophy (ingestion and digestion) of fluorescently-labeled bacteria and bacteriophages smaller than 100 nm [16]. In that study, it was estimated that the ingestion of virions might contribute up to 9% of the C, 14% of the N, and 28% of the P that phagotrophs received from ingesting bacteria. However, those estimates were considered to be underestimated due to methodological biases, and the authors suggested that under certain environmental conditions, ingesting virions might contribute similar or greater nutrition than bacteria to nanoflagellates. Furthermore, by focusing on relatively small bacteriophages, González and Suttle [16] did not address the potential full range of virions’ nutritional content, which depends on factors such as the capsid size, genome size, and presence or absence of energy-rich lipids [24, 32]. For example, the average elemental (C, N, P) content in large, dsDNA eukaryotic virus particles of the phylum *Nucleocytoviricota* [33] has been estimated to be one order of magnitude higher than in relatively small bacteriophages [32]. Additionally, these viruses contain bi-layered lipid membranes that likely increase their energy content.

In this study, we built on previous findings by quantifying the transfer of C and N (using nanoscale secondary ion mass spectrometry [34, 35]) from isotopically labeled (^13^C and ^15^N) viral particles to bacteria and protists within natural coastal microbial communities and correlated this flux to the removal of viral particles measured by flow cytometry. We performed three independent experiments with a coccolithovirus strain model system and microbial communities sampled in different years and seasons at a coastal location in the Gulf of Maine, Atlantic Ocean. Coccolithoviruses, infectious to the calcifying phytoplankton *Emiliania huxleyi* [36], were selected because of their ubiquitous distribution, ecological relevance, and morphological characteristics that represent average capsid and genome sizes of large dsDNA eukaryotic viruses. The three experiments were carried out at different times of the year but also each experimental design was improved using information obtained from the previous experiment. For example, in the first experiment, we examined the virus removal in bottles on very long timescales (up to 69 days) to get an initial idea about the turnover of viral particles with and without the microbial loop. In the second experiment, we more efficiently purified the isotope labeled viruses from algal-derived organic matter and also examined the virus- degrading microbial community structure, and in the third experiment, we separated the influence of protists from that of bacteria by adding another size fraction to the set of treatments.

## Materials and Methods

### Preparation of Emiliania huxleyi-virus (EhV) isotopically-labeled stocks

Axenic *E. huxleyi* CCMP374, originally obtained from the Bigelow Laboratory for Ocean Sciences National Center for Marine Algae and Microbiota, was maintained in seawater-based f/2-Si medium [37] (salinity ∼30 psu) at 16°C under a 14:10 h light:dark cycle at ∼100 µmol photons m^−2^s^−1^.

Fresh isotopically labeled EhV stocks were prepared prior to each experiment as follows: Exponentially growing axenic *E. huxleyi* CCMP374 were transferred (10% v/v) into sterile modified f/2-Si medium containing 3mM NaH^13^CO_3_ and 0.882 mM Na^15^NO_3_ (2.2 L for experiments 1 and 2 and 4.4 L for experiment 3, see below). The cultures were maintained in tightly closed flasks with no headspace under the standard culture conditions for about one week until they reached mid-exponential growth phase (∼0.8–1 × 10^6^ cells ml^-1^). This time was estimated based on previous characterization of *E. huxleyi* CCMP374 growth curves under similar conditions in our laboratory. Then, 5% of the culture volumes were replaced with fresh bacteria-free EhV-163 strain lysate to achieve virus:host ratios of approximately 8–10:1, and a 1 mL sample was collected and fixed with electron microscopy-grade glutaraldehyde (0.5% final concentration) for enumerating virus-like particles and confirming the absence of bacteria via flow cytometry as previously described [38]. Bacteria cells and virus-like particles were discriminated based on their side scatter signal and green fluorescence from the nucleic acid dye SYBR^TM^ Green I (Thermo Fisher). EhVs have a distinct and recognizable flow-cytometry signature that allows discrimination from other virus-like particle populations [4, 39]. The removed 5% culture volumes were kept as reference non-infected controls. The cultures were kept in the incubator until clearance was observed (3–5 days).

The lysates were filtered through 0.45 µm pore size PES membranes under sterile conditions to remove host cell debris. A 1 mL sample was analyzed by flow cytometry analysis, confirming the production of new EhV-163 particles compared to the initial sample and the absence of detectable bacteria. The lysates were then concentrated down from 2.2 L to ∼300 mL for experiments 1 and 2 and from 4.4 L to ∼150 mL for experiment 3 by tangential flow filtration (TFF) using a Vivaflow® 50, 50 KDa MWCO PES system (Sartorius) for experiment 1 and a 300 KDa MWCO QuixStand system (UFP-300-C-4MA, GE Healthcare) for experiments 2 and 3. Flow cytometry analysis of concentrate aliquots confirmed the concentration efficiency of EhV particles was 90-100% in all cases. We also confirmed by nanoSIMS analysis that the EhV particles produced with this approach were isotopically labeled (∼40% ^13^C and ∼80% ^15^N). The concentrated labeled EhV stocks were kept at 4°C until further use.

### Experiments set up and sample collection

We performed three independent incubation experiments with environmental marine microbial communities collected off the dock at Bigelow Laboratory for Ocean Sciences (Mid-Coast Maine, Gulf of Maine, Atlantic Ocean, 43° 51’ 35” N - 69° 34’ 48” W, ∼8 m and ∼11 m depth at low and high tide, respectively) supplemented with the isotopically labeled EhV-163 described above. In addition to the differences in EhV-163 stock preparations, the experiments differed in the season and year when natural communities were sampled, how those communities were size fractionated, the incubation volume, and the total duration of the experiment. The specific details are described below and summarized in supplementary Table S1.

For *experiment 1,* herein referred to as the “long incubation experiment” *(Aug. 26^th^, 2015 – Nov. 3^rd^, 2015; total 69 days)*, three liters of surface seawater were collected during incoming tide at 5:25 PM (water temperature ∼16°C, low tide: 0.84 ft, 2:20 PM; high tide: 9.76 ft, 8:42 PM). The water was gently passed through a 100 µm nylon mesh to remove most mesozooplankton and large particles, and half of that volume was subsequently filtered through a 0.2 µm pore size PES membrane using a sterile filtration unit (VWR® Vacuum Filtration Systems) (∼3 inHg vacuum pressure) to remove most cellular microbes. Triplicate 500 mL of each seawater fraction were placed in sterile borosilicate glass round bottles, and each received a 26.5 mL aliquot of the 50 KDa-concentrated EhV-163 stock (∼25 × 10^6^ EhVs mL^-1^ per bottle). Samples were collected immediately, and after 12 hours, 20 hours, 40 hours, 27 days, and 69 days of incubation at 16°C in the dark to measure particle abundances and cell isotopic labeling by nanoSIMS as follows: 20 mL aliquots from each incubation bottle were filtered onto separate 25 mm, 0.1 µm pore size polycarbonate membrane filters (Whatman® Nuclepore^TM^). The filters were rinsed with 10 mL of sterile de-ionized water to eliminate salt crystals, air dried, and stored at -80°C until analysis by nanoSIMS. Parallel 1 mL samples were fixed with glutaraldehyde (0.5% final concentration), snap frozen in liquid N_2_, and stored at -80°C until further analysis for enumerating virus-like particles and bacteria by flow cytometry following established methods [38]. Two dilutions of each biological triplicate sample were analyzed by flow cytometry. The dilutions were chosen to maintain the events per second between 100 and 500 during the analysis.

For *experiment 2*, herein referred to as the “high purification experiment” *(Jun. 29^th^, 2016 – Jul. 7^th^, 2016; total 8 days)*, eight liters of surface seawater were collected during outgoing tide at 7:50 AM (water temperature ∼15°C, high tide: 9.2 ft, 6:32 AM: low tide: -0.01 ft, 12:39 PM).

The water was passed through a 100 µm nylon mesh as in *Experiment 1*, but this time half of the <100 µm seawater was filteredthrough a 0.1 µm pore size PES membrane (VWR® Vacuum Filtration Systems) (∼3 inHg vacuum pressure) to improve the removal of small bacteria that a 0.2 µm filter might not retain. Samples for flow cytometry enumeration of virus-like particles and bacteria and for nanoSIMS analyses were collected from each seawater fraction to serve as initial controls. Additionally, we collected 200 ml of unfixed <100 µm seawater onto 47 mm 0.22 µm pore size PES filters (Millipore) for 16S rDNA and 18S rDNA sequencing. The filters were immediately stored at -80°C until analysis. Then, three 1 L borosilicate glass round bottles were filled with each seawater fraction, and 29 ml of a 300 KDa EhV-163 concentrate was added to each bottle (∼30 × 10^6^ EhVs mL^-1^). Samples were taken from each bottle immediately and after 12 hours, 24 hours, 48 hours, and 8 days of incubation at 16°C in the dark for nanoSIMS and flow cytometry analyses. In addition, on day 8, we collected samples for 16S rDNA and 18S rDNA sequencing from all replicates in the <100 µm and <0.1 µm incubations (as described above).

For *Experiment 3*, herein referred to as the “bacterial fraction experiment” *(Nov. 13^th^, 2018 – Nov. 21^st^, 2018; total 8 days)*, six liters of surface seawater were collected at 3:15 PM, approximately a half hour after high tide (water temperature ∼6°C, high tide: 8.81 ft, 2:42 PM: low tide: 0.79 ft, 9:07 PM). The water was passed through a 100 µm nylon mesh. Two liters of that seawater were sequentially filtered through 32 mm 5 µm and 1.2 µm pore size syringe filters (Pall Lab) using a peristaltic pump (150 rpm speed) to remove most protists in the 100–1.2 µm range while keeping most non-aggregated bacteria. Another 2 L of the <100 µm seawater were filtered through a 0.1 µm pore size PES membrane using a sterile filtration unit (VWR® Vacuum Filtration Systems) (∼3 inHg vacuum pressure) to produce a microbe-free seawater fraction. Six 250 ml borosilicate glass round bottles were filled with either <100 µm, <1.2 µm, or <0.1 µm seawater. Three bottles from each set received 2.1 ml of a 300 KDa EhV- 163 concentrate (∼30 × 10^6^ EhVs mL^-1^ in each bottle), while the other three were supplemented with 2.1 ml of <0.1 µm seawater to serve as no-EhV controls. Samples from each bottle were collected at the onset of the incubation and 12 hours, 24 hours, 48 hours, 4 days, and 8 days of incubation at 16°C in the dark for flow cytometry enumeration of virus-like particles and bacteria[40]. Parallel samples were collected for nanoSIMS analysis.

### Microbial Community Structure Characterization

DNA was extracted from 7 samples from the high purification experiment with the Qiagen DNEasy kit, and the 16S and 18S rRNA genes were amplified with 16S rRNA V4 primers 515F (GTGYCAGCMGCCGCGGTAA [41]) and 806R (GGACTACNVGGGTWTCTAAT [42]), and 18S rRNA V4 primers 565F (CCAGCASCYGCGGTAATTCC) and 908R (ACTTTCGTTCTTGATYRA) [43] and sequenced through the Joint Genome Institute’s Community Sequencing Program.

Paired reads (2x301 base pair) were processed with DADA2 v1.6.0 [44] for filtering (parameters maxN = 0, maxEE = c(2, 2), truncQ = 2). For the 16S reads,the forward and reverse reads were trimmed to 15-230 and 20-190, respectively. For the 18S data, both the forward and reverse reads were trimmed from 5-275. Chimeric sequences were predicted *de novo* and removed with the removeBimeraDenovo() function in DADA2 using the “consensus” method. 16S and 18S amplicon sequence variants (ASVs) were assigned taxonomy with Silva v132 [45], and the 16S ASVs were further assigned with the RDP classifier v2.11 against training set 16 [46]. The taxonomy datasets were filtered to remove ASVs with a “chloroplast” or “mitochondria” family assignment from the 16S dataset, or an “arthropoda” phylum assignment from the 18S dataset (one of the samples was contaminated with a spider 18S sequence). Data are presented as relative abundance (normalized to total read number for each sample).

### NanoSIMS and Scanning Electron Microscopy

Polycarbonate filters were thawed, air dried, and wedges of 1/8 size were cut out of each filter using sterile scissors and adhered to aluminum disks using conductive tabs (#16084-6, Ted Pella, Redding, CA) and sputter coated with ∼5 nm of gold. Isotope imaging was performed with a CAMECA NanoSIMS 50 at Lawrence Livermore National Laboratory. The primary ^133^Cs^+^ ion beam was set to 2 pA, corresponding to an approximately 150 nm diameter beam diameter at 16 keV. Rastering was performed over 20 x 20 μm analysis areas with a dwell time of 1 ms pixel^-1^ for 20-30 scans (cycles) and generated images containing 256 x 256 pixels. Sputtering equilibrium at each analysis area was achieved with an initial beam current of 90 pA to a depth of ∼60 nm. After tuning the secondary ion mass spectrometer for mass resolving power of ∼7000 (1.5x corrected), secondary electron images and quantitative secondary ion images were simultaneously collected for ^12^C_2_ ^-^, ^13^C^12^C^-^, ^12^C^14^N^-^, and ^12^C^15^N^-^ on individual electron multipliers in pulse counting mode; note that ^13^C^12^C^-^/^12^C_2_^-^ = 2 x ^13^C/^12^C and ^12^C^15^N^-^/^12^C^14^N^-^ = ^15^N/^14^N. All NanoSIMS datasets were initially processed using L’Image (http://limagesoftware.net) to perform dead time and image shift correction of ion image data before creating ^13^C/^12^C and ^15^N/^14^N ratio images, which reflected the level of ^13^C and ^15^N incorporation into biomass. To quantify substrate incorporation, biomass was identified based on ^12^C^14^N^-^ images, and regions of interest (ROIs) were drawn automatically using particle thresholding. ROIs for protists were drawn manually after identifying their location via scanning electron microscopy with a FEI Quanta model 450 scanning electron microscope with a 5 kV beam (ThermoFischer, USA). We also used an alternative approach to find protists by carrying out shallow nanoSIMS analyses over large filter areas and finding hotspots of high ^12^C^14^N^-^ signal sized from 4 to 50 microns in diameter. The isotopic ratio averages and standard error of the mean for each ROI were calculated from ratios by cycle. The ROI data were exported for further statistical analyses in GraphPad Prism. Isotope enrichment was reported either as atom percent excess, representing the percent of the normally rare isotope (^13^C or ^15^N) above natural abundance, or X_net_, the fraction of cellular component derived from the added substrate (in this case, isotope-labeled EhV, X being C or N).

### Statistical Analyses

We used a Welch’s t-test (two-sample assuming unequal variances, Alpha level 0.05) to determine the statistical significance between differences in EhV and bacteria abundances, and bacterial growth rates between incubations of different community size fractions at each sampling point, with and without the addition of EhV particles. Comparison of isotope labeling of ROIs over time were carried out using non-parametric Wilcoxon signed ranks tests.

### Food web modeling

To develop initial quantitative intuition, and initiate a framework for virovory modeling more broadly, we developed a simple foodweb model to interpret the results of the bacterial fraction experiment. The ordinary differential equation model in Equation 1 allows a population of viruses (V, ml^-1^) to be preyed upon by heterotrophic bacteria (B, mL^-1^) and a protist grazer (G, mL^1^). In Equation 1, viruses are produced at constant rate *S_v_* (viruses mL^1^ day^-1^), and in turn are fed upon by bacteria and grazers following mass action kinetics with coefficients *φ_g,v_* and *φ_b,v_* (both mL day^-1^), and decay at rate *δ_v_* (day^-1^). Bacteria grow at rate *µ_b_* (day^-1^) and consume viruses with efficiency *ɛ_b,v_* (non-dimensional, n.d.). Bacteria are in turn preyed upon by protists, also following mass action kinetics with coefficients *φ_g,b_* (mL day^-1^), and we assume bacterial mortality unrelated to protist grazing is captured with a quadratic mortality term, with coefficient *δ_b_* (day^-1^). Grazers consume viruses and bacteria with efficiency *ɛ_g,b_* and *ɛ_g,v_* respectively (both n.d.), and grazer mortality is also quadratic with coefficient *δ_g_* (day^-1^).

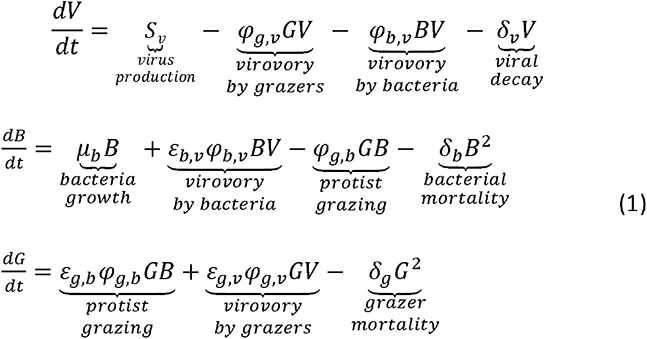

All model parameter definitions and values are provided in Table 1. The model was solved numerically with Python’s scipy odeint solver until it reached an equilibrium state (Supplementary Figure S1). Then, the bacterial fraction experiment was simulated by setting *S_v_* = 0, cutting off the supply of viruses. By running the model to equilibrium prior to simulating the experiments, we ensured that the observed dynamics were a result of the internal feedbacks prescribed in Equation 1, and not due to our chosen initial values for virus, grazer, and bacteria abundances being far from the model equilibrium state.

**Table 1:** model parameter definitions, values, and units. Parameter values were chosen to yield model solutions that were qualitatively consistent with the experimental data.

| Parameter | Symbol | Value | Units |
| --- | --- | --- | --- |
| Virus supply rate | $S_v$ | $5 \times 10^7$ | virus mL<sup>-1</sup> day<sup>-1</sup> |
| Virus consumption rate by grazer | $\varphi_{g,v}$ | $1 \times 10^{-3}$ | mL grazer<sup>-1</sup> day<sup>-1</sup> |
| Virus consumption rate by bacteria | $\varphi_{b,v}$ | $1 \times 10^{-6}$ | mL bacteria<sup>-1</sup> day<sup>-1</sup> |
| Bacteria consumption by grazer | $\varphi_{b,g}$ | $1 \times 10^{-3}$ | mL grazer<sup>-1</sup> day<sup>-1</sup> |
| Bacteria growth rate | $\mu_b$ | 0.8 | day<sup>-1</sup> |
| Virus decay rate | $\delta_v$ | 0.1 | day<sup>-1</sup> |
| Bacteria mortality rate | $\delta_b$ | $1 \times 10^{-6}$ | mL bacteria<sup>-1</sup> day<sup>-1</sup> |
| Grazer mortality rate | $\delta_g$ | 0.1 | mL grazer<sup>-1</sup> day<sup>-1</sup> |
| Transfer efficiency from bacteria to grazer | $\varepsilon_{g,b}$ | 0.1 | - |
| Transfer efficiency from virus to grazer | $\varepsilon_{g,v}$ | $1 \times 10^{-6}$ | - |
| Transfer efficiency from virus to bacteria | $\varepsilon_{b,v}$ | $1 \times 10^{-3}$ | - |

## Results

### The long incubation experiment: EhV disappearance correlated to increasing isotope enrichment of microbial biomass

In the long incubation experiment carried out in late summer, we aimed to compare the rate of EhV disappearance in flasks with and without a microbial community sampled from the coastal ocean. We note that the natural host of EhV (*E. huxleyi*) was not present or at very low background abundance in this community due to the time of the year. We found no significant reduction in EhV abundance in the flasks with either <0.2 µm or <100 µm seawater during the initial 40 hours of incubation (Fig. 1A). By the next sampling point on day 27, we measured a mean (± standard deviation) 42 ± 8 % reduction in EhV abundance in the flasks with <0.2 µm seawater, compared to the starting EhV abundances. The reduction in EhV particles in the flasks with <100 µm seawater (82 ± 9%) was significantly greater (2-sample Welch’s t-test, p = 0.0049). Thus, the presence of microbial cells roughly doubled the EhV disappearance rate from 1.7% d^-1^ to 3.3% d^-1^ (calculated between day 0 and day 27, assuming linear removal rates). By the last day of sampling (69 days), EhV abundance was further reduced to a cumulative 50 ± 7% and 96 ± 1% in the incubations with <0.2 µm and <100 µm seawater, respectively (2-sample Welch’s t-test, p = 0.0073).

**Figure 1:**
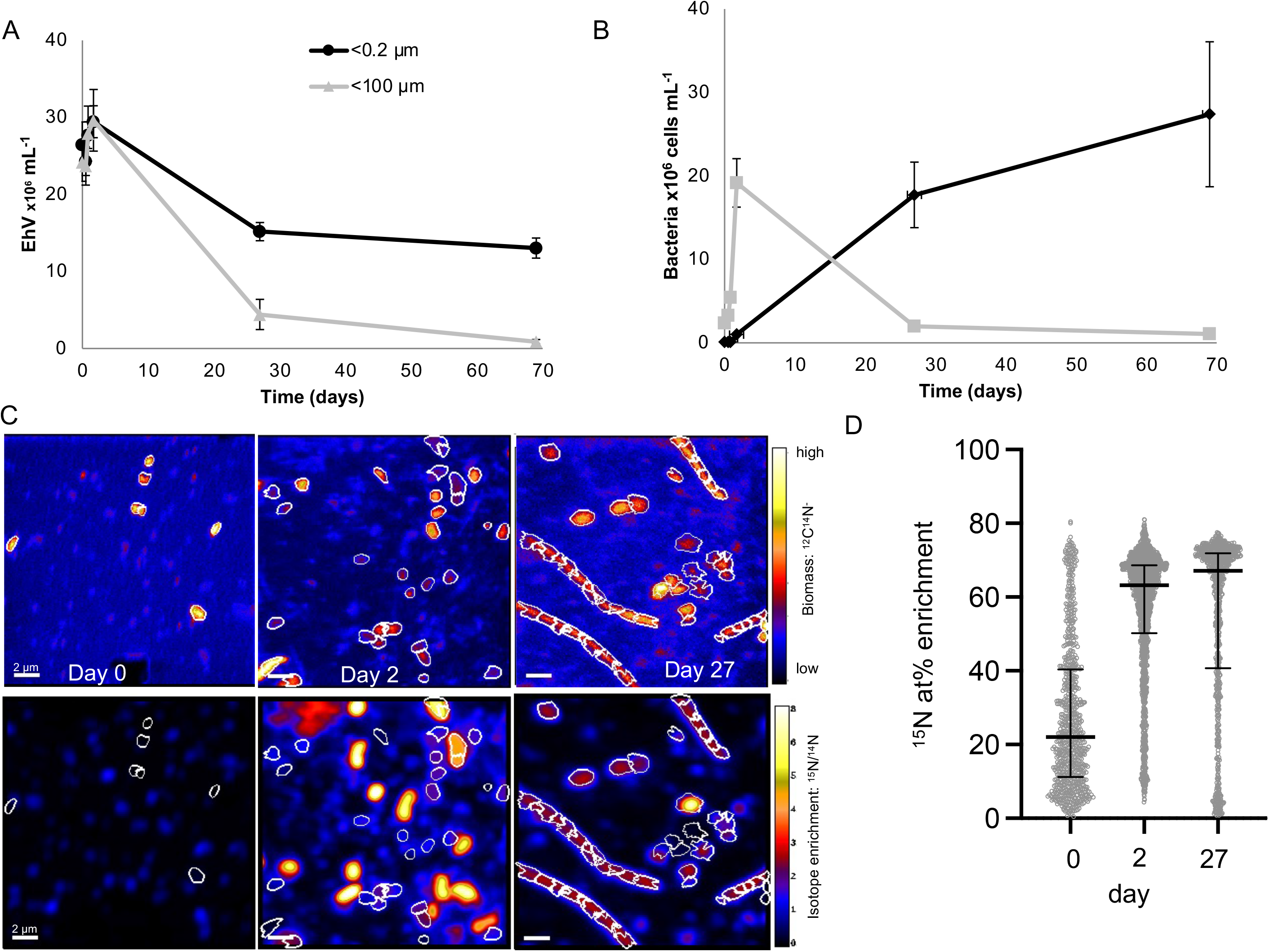
Decreases in E. huxleyi virus (EhV) over time (A) and corresponding bacterial abundances (B) as measured by flow cytometry in the long incubation experiment comparing <100 micron to <0.2 micron filtered seawater. Error bars represent ± SD. Organic particles from a representative 100 micron filtrate (identified in the top panel of C) increased in isotope enrichment over time (corresponding lower panel in C) as measured by nanoSIMS (D); scale bar = 2 μm.

Concurrent to the disappearance of EhV, we monitored the abundance of bacteria by flow cytometry (Fig. 1B). During the first 40 hours, bacteria increased in abundance in the 100 µm filtrate and as expected, were not abundant in the filtered control. As the incubations continued, however, bacteria increased in abundance in the 0.2 µm filtrate, which we attribute to the growth of a population of small bacteria that passed through the filter pores. Bacterial abundances declined in the 100 µm filtrate, likely a result of bacterivory (protist grazing on bacteria). These samples were filtered onto 0.1 µm filters and analyzed for isotopic composition by nanoSIMS (Fig. 1C). To quantify isotope incorporation by microbial cells, we first identified organic particles using the ^12^C^14^N^-^ signal as the polycarbonate filter does not contain N [47]. These ion maps show increasing numbers of organic particles from day 0 to day 2, and the appearance of long filaments by day 27 (Fig. 1C), suggesting a response of the bacterial community to a long incubation in the presence of grazers [48]. These filaments would not be identified as bacteria by flow cytometry due to their large size. We then quantified the isotopic composition of these organic particles (Figs. 1C, S2). At day zero, we found a median of 22 atom percent excess (at%) ^15^N, with a wide distribution (interquartile range of 11 to 40 at%). Isotope enrichment of particles increased after 2 days (median = 63 at%), and 27 days (median = 67 at%; Fig. 1D).

These data show a correlation between the disappearance of EhV and the isotope enrichment of the microbial loop during these long incubations and suggest that the disappearing isotopically labeled EhV were being consumed by a component of the microbial loop and their C and N became incorporated by microbial cells.

### The high purification experiment : Protists ingested EhV and incorporated viral C and N

The long incubation experiment did not examine changes between 2 days and 27 days of incubation, and the EhVs added to the incubations might have also included some high molecular weight dissolved organic matter (HMWDOM), due to the use of a relatively small molecular weight cutoff membrane (50 KDa) to concentrate the viral lysate, that likely contributed to the isotope enrichment of the microbial community. Thus, we carried out the high purification experiment (earlier in the summer of the following year) with several notable differences from the long incubation experiment. First, the EhVs were further purified from HMWDOM by using a 300 KDa filter to remove HMWDOM of 50-300 KDa that was included in the first long incubation experiment. Second, we compared the <100 µm filtered seawater to <0.1 µm filtered seawater to decrease contamination of the abiotic control with small cells that could pass through a 0.22 µm filter. Third, we sampled for a shorter duration (8 days) and collected microbial community composition data at the last sampling date to examine the eukaryotic and prokaryotic communities that were numerically dominant in the bottles with added EhV. Unlike the long incubation experiment, by day 2, EhV abundances were significantly lower in the 100 µm filtrate compared to the abiotic 0.1 µm filtrate (Fig. 2A; 2- sample Welch’s t-test, p = 0.00664) and this difference continued to increase over time. EhV mean reductions at the last sampling point (day 8) were 63 ± 9% in the <0.1 µm incubations and 91 ± 0.4% in the <100 µm incubations (2-sample Welch’s t-test, p = 0.03662). These decreased abundances correspond to abiotic and biotic removal rates of 7.8 and 11.4% d^-1^, respectively, representing a 46% increase in daily degradation rate with the presence of microbes.

**Figure 2:**
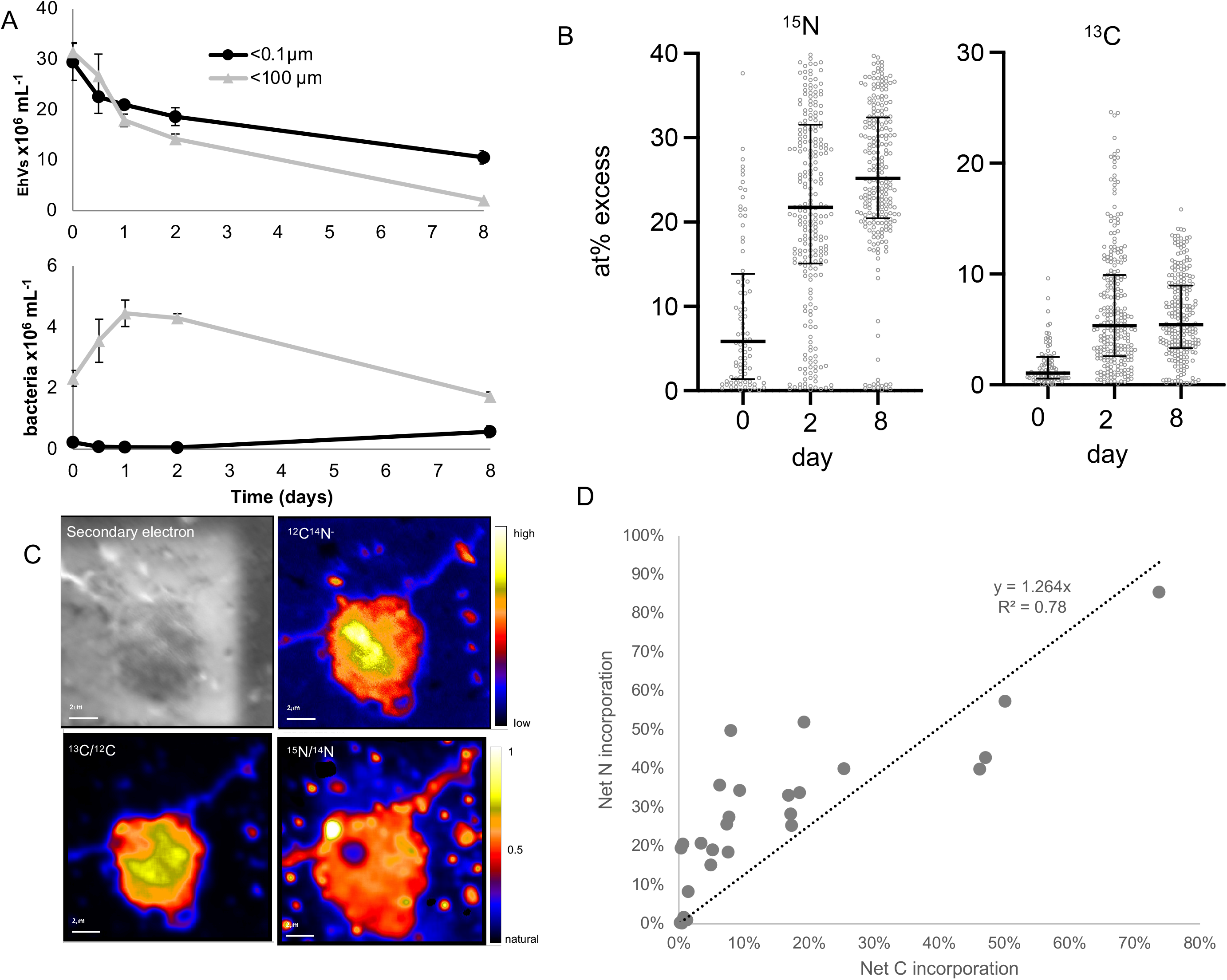
Removal of EhV and corresponding abundances of bacteria as measured by flow cytometry in the high purification experiment (A), showing increasing isotope enrichment of bacteria-sized organic particles over time (B). Individual protist cells were analyzed by nanoSIMS (one representative protist shown in panel C), and the 27 analyzed protist cells exhibited a positive correlation between ^13^C and ^15^N enrichment from the isotope labeled EhVs (D).

We again quantified the isotope composition of bacteria-sized organic particles collected on 0.1 µm filters (Fig. S3), and similarly to experiment 1, documented both increased ^15^N and ^13^C (Fig. 2B) over the length of the incubation. Isotope enrichment was lower than the long incubation experiment, likely caused by the improved purification of the viruses from HMDOM and/or batch-to-batch differences in isotope labeling of the *E. huxleyi* cultures. By day 2, median ^15^N enrichment had increased to 22 at% from 6 at% on day 0, and further increased to a median of 25 at% at day 8. Enrichment of ^13^C was lower than ^15^N but showed a similar trend. These data again show that the disappearance of isotope-labeled EhVs during the incubation was correlated with increasing isotope enrichment in bacteria-size organic particles. To directly test if protists grazed on the labeled EhV and incorporated C and N from the EhV, we used scanning electron microscopy to find protist cells from day 2 samples (Figs. 2C, S4) and analyzed those same cells (N = 27) by nanoSIMS, making sure to avoid isotope labeled hotspots near or even attached to protist cells, which likely were unincorporated EhV (Fig. S4). To quantify the contribution of virus N and C to protist N and C demand, we calculated N_net_ and C_net_, the percent of biomass N and C that originated from EhV N and C [49]. Note that N_net_ or C_net_ values of 50% indicate one cellular division, where 50% of the cell’s biomass originated from the added substrate (assuming no other source of C or N), and two divisions correspond to C_net_ or N_net_ values of 75%. C_net_ and N_net_ were positively correlated and ranged from 0.2% to 85% (Fig. 2D). We interpret the positive correlation as evidence for direct incorporation of virus C and N rather than cross-feeding from remineralized N, where we might expect only ^15^N enrichment. We also do not know the extent of mixotrophic EhV ingestion since our incubations were carried out in the dark, inhibiting photosynthesis. These data directly show that a subset of protists intensely grazed on EhV during at least the first 2 days of incubation and incorporated, in some cases, very high levels of C and N, corresponding to multiple cell divisions using viral C and N as the sole source of nutrition.

We collected small subunit ribosomal DNA sequences from day 8 samples and compared them to day zero to examine the microbial (bacterial and protistan) community changes with added EhV (Fig. 3). As expected, there were very few eukaryotic sequences (18S) in the 0.1 µm filtrate incubations; the few reads in those samples likely originated from amplification of environmental DNA and we did not analyze these data any further.

**Figure 3:**
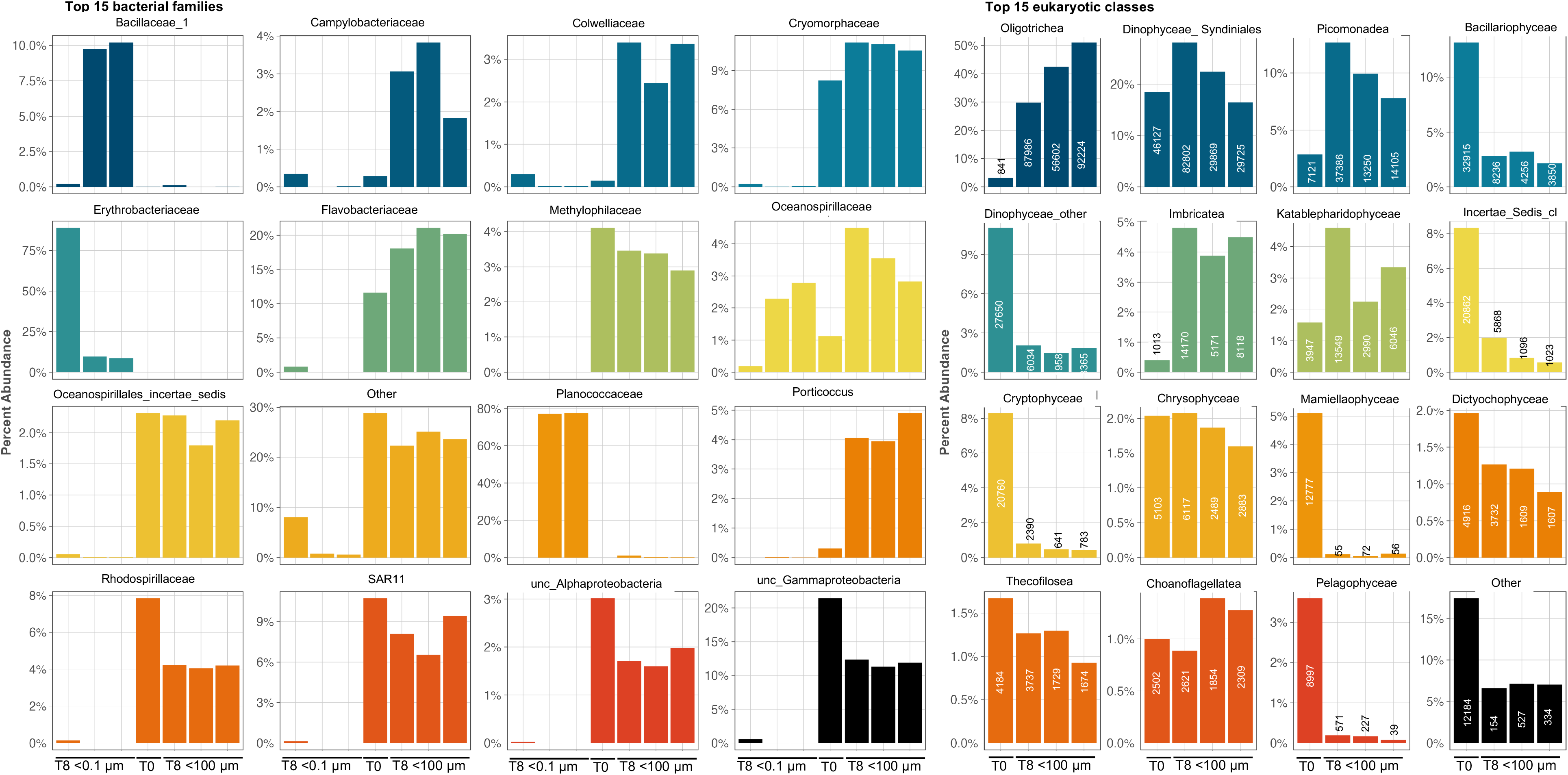
Relative abundance of 16S rRNA bacterial families (left) and 18S rRNA eukaryotic classes (right, including the number of reads detected) from the high purification experiment collected at the end of the 8-day incubation, comparing the replicate <0.1 and/or <100 micron filtrates to the community at inoculation (T0).

On day 8, the microbial communities in the 0.1 µm filtrate incubations were dominated by bacteria from the families *Erythrobacteraceae* (>75% relative abundance in one biological replicate), or *Bacillaceae* and *Planococcaceae* (∼10% and ∼80%, respectively, in the other two biological replicates). These bacterial families were absent at T0 in the 100 µm filtrate environmental water and absent or negligibly represented in the bacterial communities within the 100 µm filtrate incubations at T8. *Oceanospirillaceae* reads found in intermediate abundance in the 0.1 µm filtrate incubations (<3% relative abundance), were also present at intermediate abundance in the 100 µm filtrate incubations. Notably, *Erythrobacteraceae* had opposite abundance trends to *Bacillaceae*, *Planococcaceae*, and *Oceanospirillaceae*. Members from the families *Planococcaceae* and *Bacillaceae* form spores [50], which could explain how those bacteria entered and developed in the the 0.1 µm filtrate incubations. The families *Oceanospirillaceae* and *Erythrobacteraceae* include members with ultra small cell sizes that likely pass through 0.1 µm pore-size filters [51]. It has also been shown that some marine bacteria can pass through filters and recover to larger sizes after some time allowed for growth [52]. The <100 µm filtrate flasks were dominated by bacterial taxa absent or not significantly represented in the 0.1 µm filtered samples. These taxa included families commonly identified in surface marine samples [53]. The detected bacterial families responded differently to the incubation with EhVs. Some largely increased their relative abundance, either through growth or decline of other taxa by T8 (*Campylobacteracea*, *Colwelliaceae*, *Oceanospirillaceae*, *Porticoccaceae*), others had a moderate increase (*Cryomorphaceae*, *Flavobacteriaceae*) and other taxa declined or had little relative abundance chances between T0 and T8 (e.g., *Methylophilaceae*, *Pelagibacteriacea*, *Rhodospirillaceae*).

Assumed photosynthetic protists (e.g., Bacillariophyceae, Prasinophytae, Dinophyceae, Cryptophyceae, Mamiellaophyceae, and Pelagophyceae) that were relatively abundant at T0 had a drastic decline by T8, as expected since the incubations with EhV were carried out in the dark. Some taxa did not change in relative abundance between T0 and T8, such as the Chrysophyceae, Syndiniales (a subset of dinoflagellates), Dictyochophyceae, Thecofilosea (amoebae), Katablepharidophyceae (cryptophytes), and Choanoflagellatea. Eukaryotic taxa with high relative abundance on the last day of the incubations (defined as having > 8,000 18S rRNA reads from at least 2 of the 3 replicates) included the heterotrophs Oligotrichea (ciliates), Picomonadidea (picobiliphyte), and Imbricatea (cercozoa) (Fig. 3). Furthermore, these taxa showed a large increase in relative abundance (and number of reads) between T0 and T8 in the three incubation replicates. Consequently, these taxa have a high likelihood of representing the protists that grazed the added EhV. Oligotrich ciliates are known to be generalist grazers, but interestingly, picobiliphytes have been identified as heterotrophs that obligately consume small particles < 150 nm, the size of EhV [54]. Furthermore, a recent study using single cell genomics showed convincing evidence that a number of protistan lineages, including picobiliphytes, within a water sample collected nearby our sampling site in the coastal Gulf of Maine likely consumed viruses [19]. Our nanoSIMS data also identified isotopically labeled protist cells of various sizes and shapes, which together with the eukaryotic taxonomy data, suggest that multiple taxonomic groups grazed on EhV in the incubations.

### The bacterial fraction experiment: bacteria removed EhV and incorporated viral C and N without the presence of protists

The first two experiments (“long incubation” and “high purification”) examining EhV removal compared a <100 µm microbial community to one with most cells removed (<0.1 or <0.2 µm) and quantified the incorporation of EhV-derived C and N into protist and bacteria cells. In the third “bacterial fraction” experiment (carried out in mid-fall), we aimed to disentangle the effect of protists and bacteria by comparing EhV disappearance across 3 size fractions: <100 µm (included protists and bacteria), <1.2 µm (bacteria-enriched fraction), and <0.1 µm (most cells removed). We also monitored the abundance of bacteria in side-by-side incubations without the addition of EhV to quantify how much bacterial biomass was generated from the addition of EhV as a nutrition source, and we also added another sampling time at day 4. After 4 days, the <100 µm incubation had significantly lower EhV abundances than the <0.1 and <1.2 µm treatments (2-sample Welch’s t-test, p = 0.00005; Fig. 4A). After 8 days however, both the <1.2 and <100 µm treatments had significantly less EhV than the <0.1 µm treatment (2-sample Welch’s t-test, 0.1 µm vs. 100 µm p = 0.013; 0.1 µm vs. 1.2 µm p = 0.0011), corresponding to mean reductions of 60 ± 7.0% and 83 ± 1.5%, respectively, compared to reductions of 45 ± 7.6% in the <0.1 µm flasks. We calculated daily removal rates to be 5.6%, 7.5%, and 10.4% d^-1^ in the <0.1 µm, bacteria-enriched, and bacteria and protists flasks, respectively. This corresponds to a 33% increase in removal rate with bacteria and 84% increase with protists compared to EhV removal in the flasks that had most cells initially removed by filtration.

**Figure 4:**
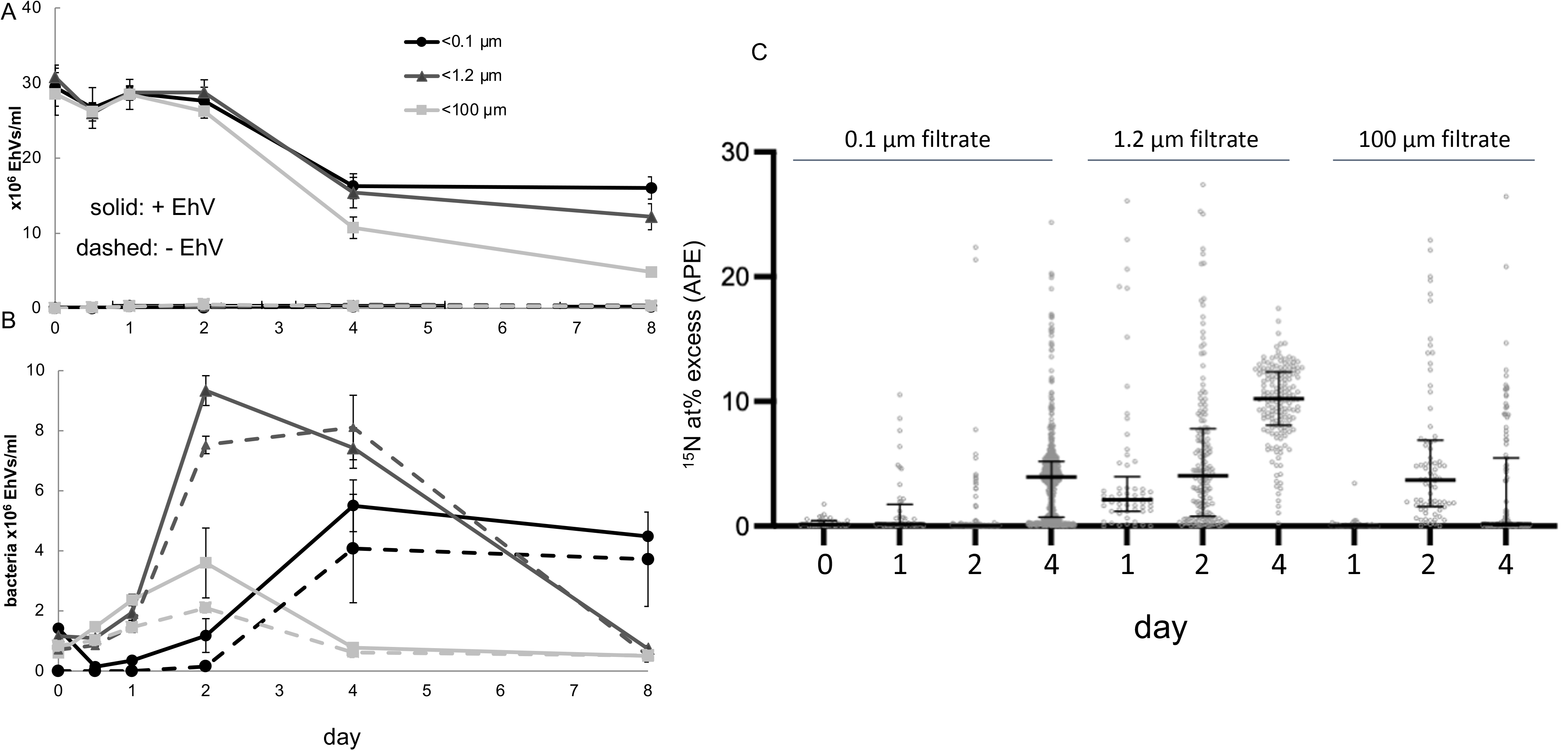
Removal of EhV (A) and corresponding bacterial abundances (B) from the bacterial fraction experiment, showing some EhV removal without protists. The nanoSIMS data confirmed that bacteria in the <1.2 micron treatment became increasingly isotope labeled over time (C).

Bacterial abundances in the <1.2 and < 100 µm treatments, with or without EhV added, were not statistically different within the first 24 hours (Fig. 4B). However, by day 2 bacteria grew to higher abundances with EhV added compared to without, both in the <100 and <1.2 µm treatments (2-samples Welch’s t-test, p<0.001). This suggests that EhV C and N supported bacterial growth, but there was a 48 hour lag until this increase was noticeable. We also note that the <0.1 µm treatment became contaminated with bacteria earlier than the first two experiments, and bacterial abundances in these flasks became higher than the <100 µm flasks as soon as day 4.

Similar to the long incubation and the high purification experiments, the increase of bacterial abundances in both the bacteria-enriched and the full microbial community was associated with increases in ^15^N enrichment of particles over time (Fig. 4C). First, we examined particles in the <0.1 µm treatment and found no increase in isotope labeling through day 2, up to a median of 3.9% APE ^15^N by day 4 (Fig. 4C). On the other hand, the bacteria-enriched treatment, which likely included some very small protists, showed increasing labeling on days 1, 2, and 4, consistent with bacterial degradation and incorporation of EhV N, this time without the presence of protist grazing. Interestingly, the bacteria-sized particles with the protists showed increasing isotope labeling by day 2 but then decreasing down to background isotope composition by day 4 (Fig. 4C). This suggests that the protists were likely grazing on both virions and bacteria (both of which decreased between day 2 and day 4), which removed isotope-labeled particles and leaving non-growing (and mostly unlabeled) material, including cells.

### Food-web modeling to capture virus removal dynamics

We next assessed the ability of a simple food-web model to capture the dynamics, measured by flow cytometry, for viruses and bacteria in the bacterial fraction experiment (Fig. 5). To reproduce filtration by 100, 1.2, and 0.1 µm, we assumed the sequential removal of all microzooplankton larger than 100 µm, removal of protists, and removal of bacteria, respectively. The model captured diminishing removal of viruses with increasingly fine levels of filtration (Fig. 5a), as documented in the experiment. It also captured the rise in bacteria with removal of protists in both the 1.2, and 0.1 µm treatments (Fig. 5b). The model also predicted a modest decline in protist grazer abundance in the <100 µm due to the nutritional shortfall associated with loss of viruses (Fig. 5c). These results shows that relatively simple simulations can recapitulate our experimental data, but this food-web model has limitations preventing its application to natural ecosystems. For example, there is only one group of viruses, when in practice there are diverse assemblages of viruses that will respond differently to consumption by protists and bacteria. Furthermore, we did not account for changes in protist losses associated with the removal of microzooplankton, and the potential this could have to modify bacteria dynamics. Nevertheless, the gradient in mortality associated with filtration (Fig. 5a) provides a basis on which to develop more realistic models of virovory within a food-web context in the future.

**Figure 5:**
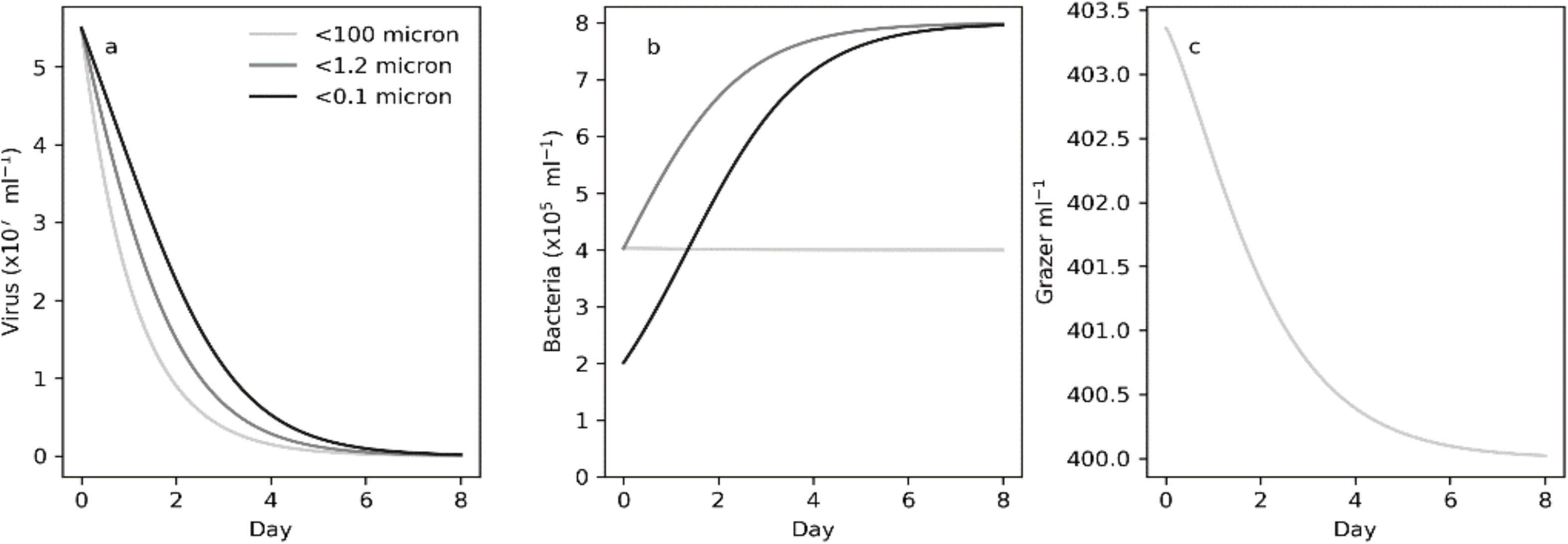
Food web modeling results that recapitulated experimentally measured virus (a) and bacterial (b) abundances based on including protist and bacterially-control virus decay and protist grazing of bacteria. Also assumed was constant protist mortality over time in the long bottle incubations (c).

## Discussion

Here, using laboratory flasks experiments, we documented the removal of viral particles and the concurrent transfer of viral C and N into bacteria and protist biomass. Given slightly different alterations to the experimental designs and the time of year, the data were relatively consistent, showing that without cells (or with most of them initially filtered out), viral removal reached 45 to 63% within one week, and the presence of the microbial loop increased this removal from 83 to 96%. It is unknown if the 0.2 or 0.1 μm filtrates represented over or under-estimates of abiotic viral particle removal. The flasks were incubated statically (without shaking) which would decrease particle aggregation caused by shear forces, which we might expect to underestimate virion removal. However, the high surface area to volume ratio of the relatively small laboratory flasks might increase particles sticking to the flask walls, which might overestimate virion removal.

Our ability to quantify the transfer of viral C and N into different types of cells (bacteria and protists) was dependent upon the high spatial resolution of the nanoSIMS instrument (∼ 100 nm). This allowed us to target regions of the filters that were either identified as protist cells via SEM, or as regions of bacteria-sized organic material that might include bacterial cells and particulate detritus. However, the nanoSIMS still was not able to image single EhV particles because they are roughly the size of the beam, and thus single particles on the images could have been 2 or more EhV particles stuck to one another. However, the resolution still enabled us to identify if protist cells were truly isotope enriched or rather had isotope labeled virions stuck to the outside of the cells (see supplementary figure S5 as an example). Also, we noticed that for some of the experiments, larger particle regions of interests at day 0 (right after inoculation) were often highly isotope enriched, suggesting that the virus particles aggregated unto themselves or other organic particles. Thus, it was crucial for us to quantify isotope enrichment of particles over time rather than sampling only one time point.

Modelers are beginning to resolve the impacts of viruses on ocean ecology and biogeochemistry [56–58]. However, the impact of virovory on global cycles of C, N, and P is, to the best of our knowledge, completely unexplored in biogeochemical models. Our foodweb model provides an initial framework for resolving this pathway in models, but there are many limitations. The parameters were chosen to illustrate the ability of the model to capture the experimental dynamics, but are not quantiatively accurate. Future work could attempt to constrain model parameters by measuring protist grazer abundances across treatments and then fitting the model to a time-series for virus, host, bacteria, and protist abundances.

Quantification of model parameters would allow direct incorporation of virovory within ocean ecological and biogeochemical models. Moreover, we also did not examine the impact of viral concentration on removal rate (we added the same concentration of viruses to the 3 experiments and did not vary it). Viral concentration is likely to have a strong influence on the microbial community response. The abundance of viral particles we added to the incubations were typical of abundances found during mesocosm-induced algal blooms infected by viruses [59], so is on the higher end of what might be expected in a natural ecosystem [60, 61]. Future experiments should examine the impact of viral concentration, including natural abundances across space and time, on their removal rates.

## Acknowledgements

Initial seed funding for this work was provided by the Gordon and Betty Moore Foundation’s Marine Microbiology Initiative projects # GBMF4465 & Charitable Contribution for Bigelow Laboratory for Ocean Sciences #4465, and work was subsequently supported by the US Department of Energy’s (DOE) Genomic Science Program through the LLNL μBioSpheres Science Focus Area grant # SCW1039. Part of this research was carried out at Lawrence Livermore National Laboratory (LLNL) under Contract DE-AC52-07NA27344. Ribosomal RNA sequencing was carried out at the Joint Genome Institute (JGI) through Community Sequencing Program award # 1939. JGI is sponsored by the DOE’s Biological and Environmental Research program and operated under Contract Nos. DE-AC02-05CH11231.

## Notes

### Competing Interest Statement

The authors have declared no competing interest.

