## Supplementary table and figures for "Coastal bacteria and protists assimilate viral carbon and nitrogen"

**Supplementary material for “Coastal bacteria and protists assimilate viral carbon and nitrogen”**

Table S1 Experimental design summary.

| | EhV lysate purification | EhV lysate concentration | Microbial community sampling time | Microbial community size fractions | Incubation volume | EhV addition | Sampling time points | Analyses |
|---|---|---|---|---|---|---|---|---|
| **Experiment 1** | <0.45 µm<br>50 KDa | 2.2 L to 300 ml | Late summer<br>Incoming tide | <100 µm<br><0.2 µm | 500 ml | ~25 × $10^6$ EhV $ml^{-1}$ | 0 h, 12 h, 20 h, 40 h, 27 d, 69 d | flow cytometry (virus and bacteria abundance) nanoSIMS (isotope labeling) |
| **Experiment 2** | <0.45 µm<br>300 KDa | 2.2 L to 300 ml | Early summer<br>Outgoing tide | <100 µm<br><0.1 µm | 1000 ml | ~30 × $10^6$ EhV $ml^{-1}$ | 0 h, 12 h, 24 h, 48 h, 8 d | flow cytometry (virus and bacteria abundance) nanoSIMS (isotope labeling) amplicon sequening (16S rRNA + 18SrRNA) |
| **Experiment 3** | <0.45 µm<br>300 KDa | 4.4 L to 150 ml | Late autumn<br>~high tide | <100 µm<br><1.2 µm<br><0.1 µm | 250 ml | ~30 × $10^6$ EhV $ml^{-1}$ | 0 h, 12 h, 24 h, 48 h, 4 d, 8 d | flow cytometry (virus and bacteria abundance) nanoSIMS (isotope labeling) |

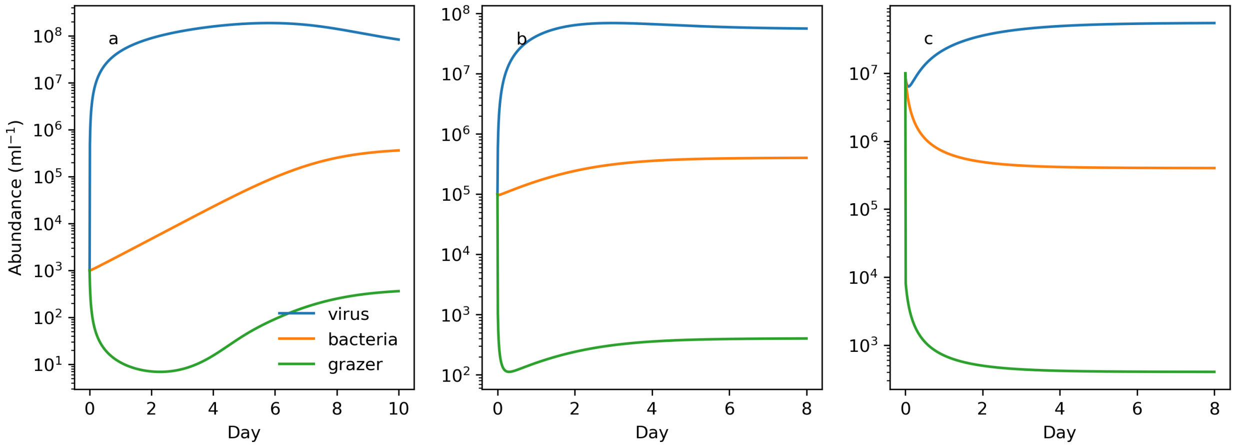


*Figure S1: Food web model preliminary simulation ('spin-up') allowing the system to reach equilibrium prior to simulating the bacterial fraction experiment. All model parameter values are provided in Table 1 of the main text. The model converged to the same set of equibrium values regardless of initial conditions, as demonstrated with sensitivities assuming initial abundances for viruses, grazers, and bacteria were a) 1x10$^{3}$, b) 1x10$^{5}$, and c) 1x10$^{7}$ mL$^{-1}$.*

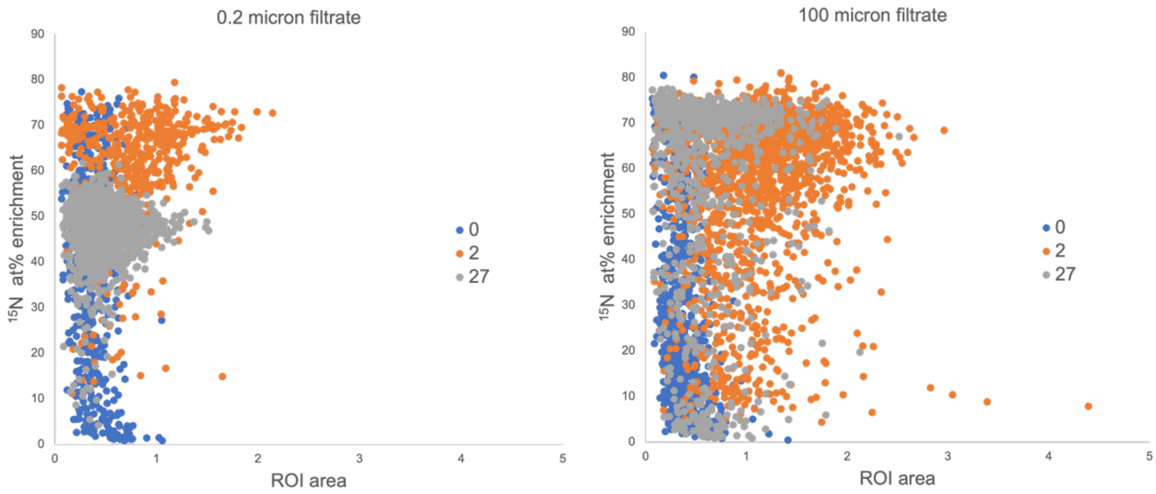


Figure S2: $^{15}N$ isotopic enrichment of regions of interests (ROIs) of different sizes from the long incubation experiment, collected from nanoSIMS images. ROIs were drawn automatically using the $^{12}C^{14}N^{-}$ images (see figure 1).

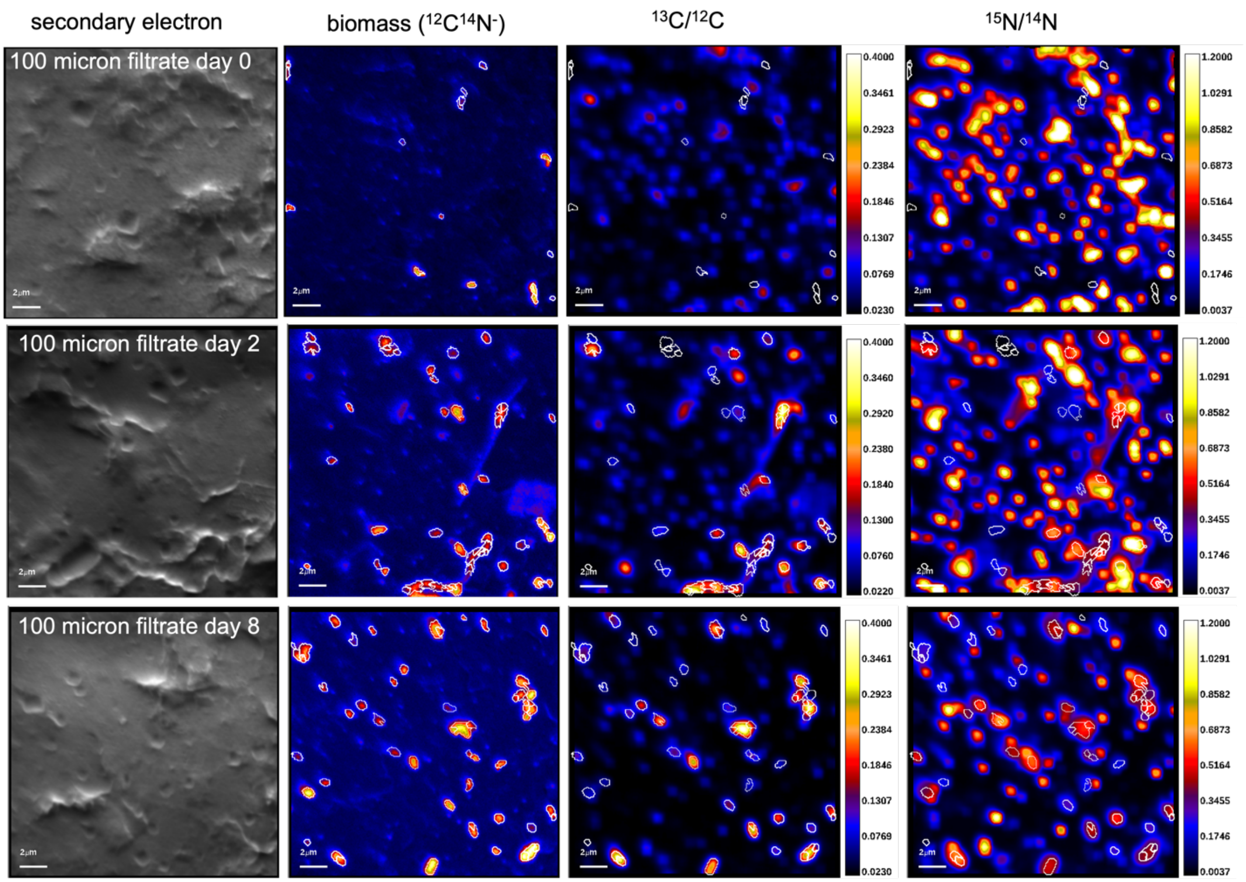


Figure S3: representative nanoSIMS images from the high purification experiment. Regions of interests (ROIs, circled in white) for isotope quantification were chosen based on the $^{12}C^{14}N^{-}$ images that identify organic particles (the filter does not contain N). On day 0, the isotope labeled particles that represent the labeled EhVs generally do not correspond to those ROIs. After 2 and 8 days, some of the ROIs are labeled, some are not, and there still remain many isotope labeled particles that do not correspond to ROIs (representing remaining EhVs).

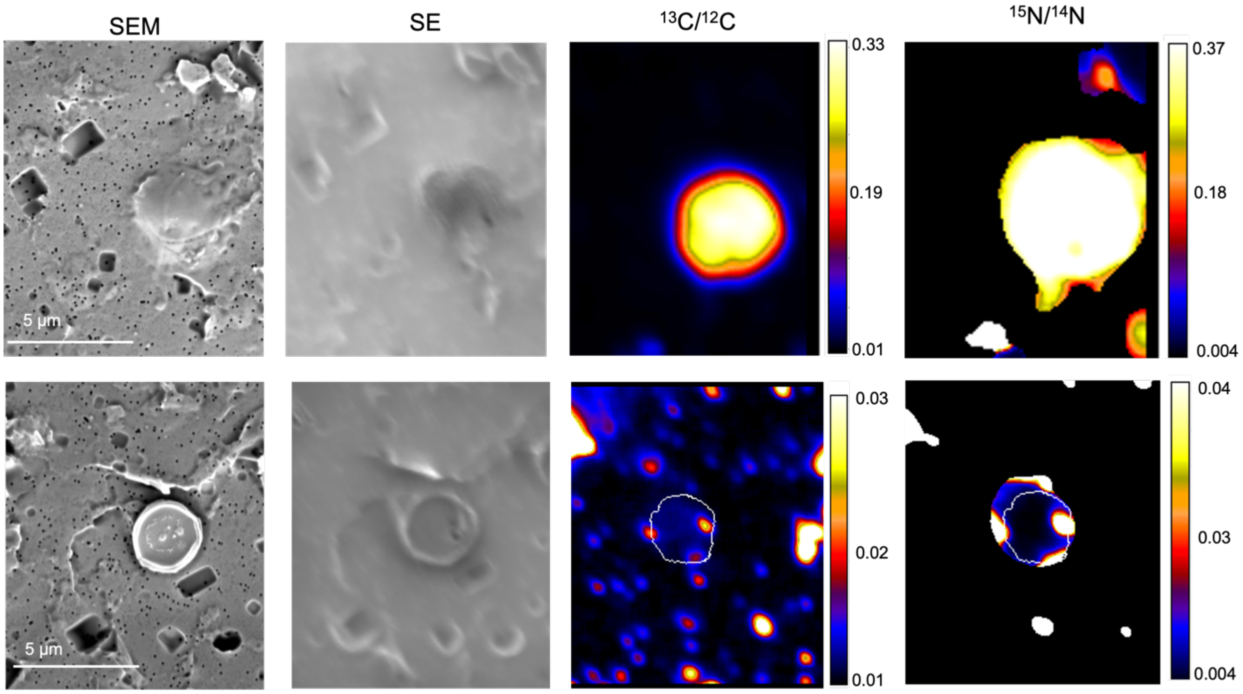


Figure S4: SEM and nanoSIMS images from protists identified in the high purification experiment sampled on day 2. The protist shown in the top panel is highly labeled in $^{13}C$ and $^{15}N$. The protist shown on the bottom is not isotopically enriched (the isotopically enriched hot spots next to the cell are likely viral particles or bacterial cells that became attached or were filtered underneath the protist cell.
